# Early life social isolation primes flies for more rapid TDP-43 dependent neurodegeneration later in life

**DOI:** 10.64898/2026.09.23.753732

**Authors:** S. MurthyGowda, K. Huyghue, B. Castillo, F. Gugala, W. Li, J. Dubnau

## Abstract

A key pathophysiological hallmark of amyotrophic lateral sclerosis (ALS) and frontotemporal dementia (FTD) is loss of nuclear localization and abnormal cytoplasmic aggregation of TAR DNA-binding protein 43 (TDP-43). Upstream factors that trigger the onset and progression of such neurodegenerative diseases are largely unknown. Aging and environmental factors contribute as potential risk factors. But epidemiological evidence suggests psychological stressors such as social isolation and loneliness are also associated with increased risks of neurodegenerative disease [1–8]. We examined the impacts of social isolation on TDP-43 related neurodegeneration. To investigate the potential for causative impact on neurodegeneration, we compared the effects of social isolation versus social enrichment in *Drosophila*. We utilized an established social isolation paradigm in which flies were either housed alone or in groups as young adults. We found that early-life social isolation acts as a primer to exacerbate the rate of neurodegeneration in response to subsequent induction of pathological levels of TDP-43. Social isolation stress drives more rapid activation of mdg4, an endogenous retrovirus that functionally mediates TDP-43 effects and more aggressive propagation of TDP-43 protein pathology from surface glial cells to nearby neurons. Shortened life span also ensues. We demonstrate that providing isolated flies with visual, olfactory and tactile social cues from conspecifics living behind a divider was insufficient to alleviate these effects of social isolation on neurodegeneration. Our findings have implications for the association between psychological stressors such as loneliness and risk of neurodegenerative diseases in humans and provide a platform to investigate mechanistic underpinnings in animal models.

## Introduction

Social isolation and loneliness are risk factors for diagnosis of neurodegenerative disorders including AD, FTD and PD [4–12]. But it is not clear whether the association reflects a causative relationship. A key pathophysiological hallmark for ALS, FTD and AD is the loss of nuclear localization and abnormal cytoplasmic aggregation of TDP-43 [13, 14]. Such pathology is observed in the affected brain regions of ∼95% of ALS, ∼40% FTD and many cases of AD [14–20]. The disruptive cellular consequences of the nuclear TDP-43 loss and accumulation of cytoplasmic inclusions are diverse, but convergent findings from postmortem tissue studies, animal models of disease and cell culture experiments include broad effects on splicing [21–23], dysfunctional nuclear-cytoplasmic transport [24–26] and disruption of mechanisms to silence retrotransposons and endogenous retroviruses (RTEs and ERVs) [27–40].

Mutations that alter the amino acid sequence of TDP-43 cause a small fraction of ALS cases, but most ALS, FTD and AD cases are sporadic, with no known genetic causes. Thus, most cases where TDP-43 protein pathology is seen contain inclusions of TDP-43 with wild type amino acid sequence. Upstream factors that trigger onset and modulate the rate of progression of neurodegenerative disease are incompletely understood, but there is growing evidence that cellular features of the normal aging process and environmental stressors each contribute [41–51]. Neuropsychiatric diagnoses such as anxiety and depression or post-traumatic stress disorder (PTSD) as well as psychological stressors such as loneliness are each implicated as risk factors that are associated with increased rates of subsequent diagnosis with neurodegenerative disease [1–3].

To investigate the potential for causative impact on neurodegeneration, we compared the effects of social isolation versus social enrichment in *Drosophila*. We utilized an established social isolation paradigm in which flies were either housed in isolation or in groups as young adults. We found that early-life social isolation acts as a primer to exacerbate the rate of progression neurodegeneration in response to subsequent induction of pathological levels of TDP-43. We found that prior social isolation stress exacerbates the effects of subsequently induced TDP-43 pathology on lifespan, drives more aggressive non-cell autonomous effects of glial TDP-43 pathology on nearby neurons and results in more rapid activation of mdg4-ERV. We also demonstrate that the protective effect of group housing requires a holistically complete social interaction that requires that animals share the same physical environment.

## Results

### Social Isolation stress exacerbates effects on lifespan from subsequent induction of pathological levels of TDP-43 in surface glia

Social isolation is highly stressful to most organisms, including *Drosophila* [52–66]. In flies, chronic social isolation causes changes in sleep, memory, feeding, metabolism and proteostasis [9, 67–77]. We made use of a chronic social isolation paradigm **(Fig 1A)** that is known to cause daytime sleep loss in flies [9], to test for effects on TDP-43-mediated neurodegeneration. Social isolation was achieved by placing newly eclosed male flies alone in a standard food vial. Social enrichment was achieved by placing newly eclosed males in groups of 20 in standard food vials. After varying lengths of such isolated or grouped housing, the male animals were placed into fresh food vials in groups of 10 (**Fig 1 A-G**) to measure their lifespans. To test the effects on neurodegeneration, we used the established Gal80^ts^ approach to induce cell type specific over-expression of human TDP-43 (hTDP-43) in either all glial cells (Repo-Gal4) or in the subperineural glia (SPG-Gal4), which help form the blood brain barrier. We have previously established that induced expression of human TDP-43 with either of these glial driver lines is sufficient to trigger aggregation pathology, loss of nuclear localization, and DNA damage within the target glial cell population and to cause a spreading effect leading to aggregation of TDP-43, DNA damage and cell death in nearby neurons that express physiological levels of either the endogenous fly ortholog of TDP-43 or of human TDP-43 [33, 34].

**Fig 1:**
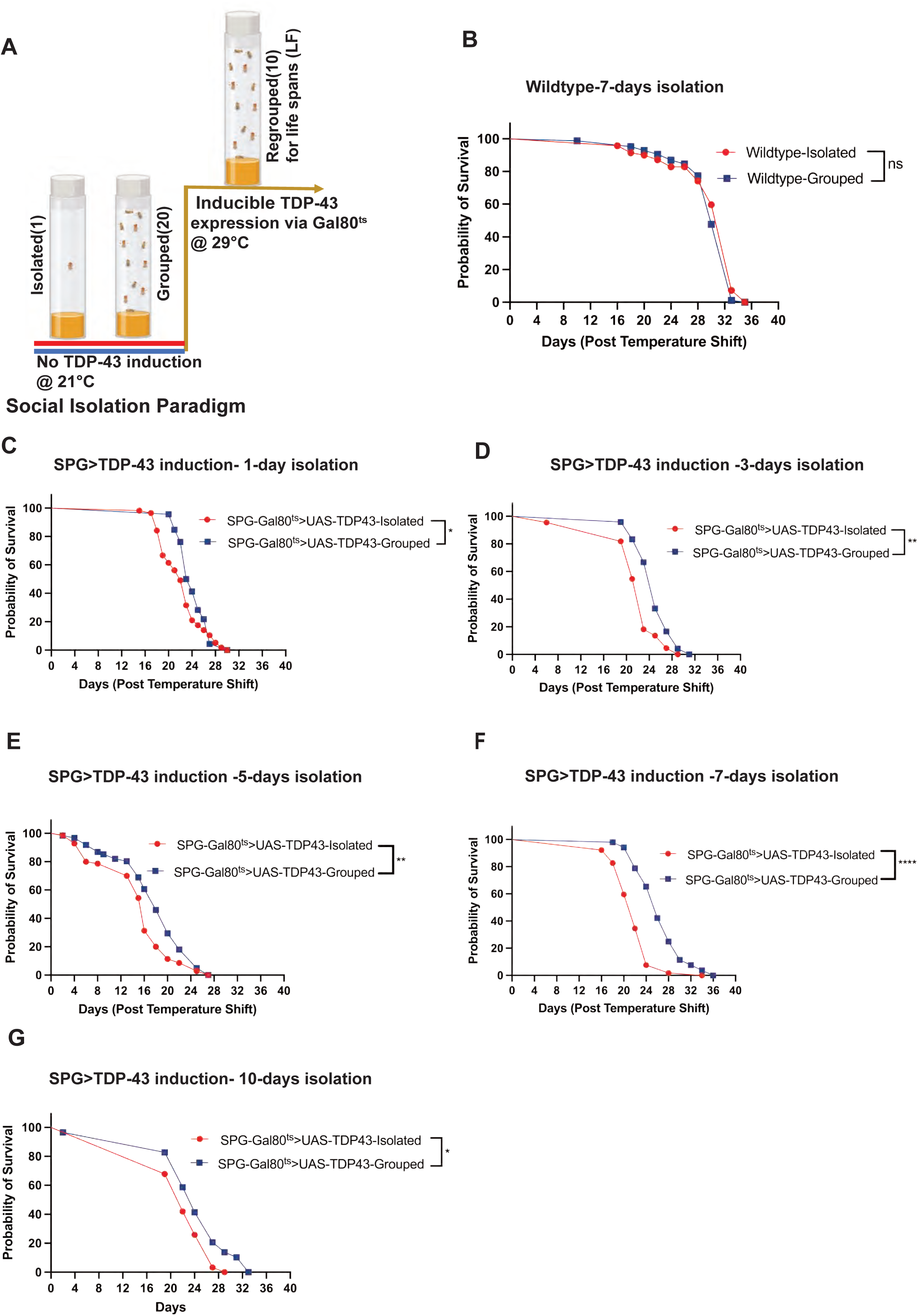
Social isolation during early adult life causes shorter lifespan in response to subsequent induction of TDP-43 pathology in SPG. **A)** Immediately after eclosion, adult male flies were housed in groups of 20 or isolated individually for varying numbers of days at 21°C prior to TDP-43 induction at 29°C. After the period of grouped or isolated housing at 21°C they were placed in groups of 10 per vial and shifted to 29°C for TDP-43 induction in SPG (surface glial cells). Temperature control of expression was achieved via the Gal80^ts^ system. Lifespans were measured at 29°C. **B)** 7 days of early life social isolation has no effect on subsequent lifespan of wild type males. Median survival after such social isolation in male flies was 33 days (n=70) vs 30 (n=84) after grouped housing. ns., not significant (Log-rank test). **C-G)** By contrast, early life social isolation for 1, 3, 5, 7 or 10 days primes flies for more rapid decline when TDP-43 is induced in SPG after the isolation period. Median survival with SPG>TDP-43 induction after 1 day of isolated vs grouped housing (C) was 22 (n=57) vs 23.5 (n=46). *p<0.05 (Log-rank test). Median survival after 3-days of isolated vs grouped housing was 23 (n=22) vs 25 (n=24). **p<0.01 (Log-rank test). Median survival after 5 days of isolated vs grouped housing (**E)** was 16 (n=70) vs grouped 18 (n=61). **p<0.01 (Log-rank test). Median survival after 7 days of isolated vs grouped housing (**F)** was 22 (n=52) vs 26 (n=52). ****p<0.0001 (Log-rank test). And median survival after 10 days of isolated vs grouped housing (**G)** was 22 (n=31) vs 24 (n=29). *p<0.05 (Log-rank test). Survival curves are shown for SPG-Gal80^ts^>UAS-TDP-43. Simple survival analyses-Kaplan-Meier –method was performed for all survival plots.

We first tested the impact of chronic (7 day) early life social isolation stress on the lifespan of wild-type adult *Drosophila*. To match experimental conditions used to induce TDP-43 overexpression via the Gal80ts system (see below), we conducted the early life social isolation vs grouped housing of wild type males at 21°C and the lifespan analyses were conducted subsequently with 10 males grouped at 29°C **(Fig 1A)**. We found that 7 days of such early life social isolation did not impact the longevity of wild type control flies (**Fig 1B**). We next tested the effects of 1, 3, 5, 7 or 10 days the same social isolation vs group housing conditions on flies that contained the SPG-Gal4, the UAS-hTDP-43 and the Gal80ts transgenes (SPG-Gal80ts>UAS-TDP-43). Animals with this combination of transgenes allow for SPG cell type specific induction of TDP-43 contingent upon switching the temperature from 21°C to 29°C. Male flies of this genotype were either socially isolated or housed in groups of 20 at 21°C to maintain Gal80 repression of the UAS-TDP-43 transgene. After this period of group vs isolated housing, the males were maintained in groups of 10 and shifted to 29°C to induce over-expression of hTDP-43 in the SPG glial cells. Flies were then maintained at 29°C for the duration of the lifespan analyses **(Fig 1A)**. We found that socially isolating SPG-Gal80^ts^/UAS-TDP-43 males (SPG^ts^>TDP-43) for either 1, 3, 5, 7 or 10 days was sufficient to prime the animals to become more sensitive to effects of subsequent induction of TDP-43 pathology because it significantly decreases median lifespans compared to their group housed counterparts (**Fig 1C-G**). We also tested the effects of prior social isolation on survival in response to subsequent TDP-43 induction in all glia using the Repo-Gal4 line, again in combination with the Gal80^ts^ and UAS-hTDP-43 transgenes (**S1A Fig**). In this case, we do not see effects of social isolation, likely due to the already rapid timeline with this more aggressive neurodegeneration model. Indeed, the median survival was 14 days for grouped flies with such pan-glial induction of TDP-43, about half that observed with induced expression restricted to the SPG, with median lifespan of 26 days. The priming effect of social isolation on response to TDP-43 pathology in SPG is sex specific because 7-days of isolation does not impact survival in SPG^ts^>TDP-43 females under identical conditions (**S1B Fig**). We also compared the effects on males of 3, 7 or 10 days of social isolation versus group housing with other males vs with females. The protective effects of group housing on males are largely similar irrespective of the sex of the conspecifics that they are housed with (**S1 C-E Fig**).

Because females are resilient to these effects of social isolation, we focused on the effects of social isolation on males. To avoid the potentially complex biological effects of courtship and reproduction, we also focused for the remaining experiments on a comparison of males that experienced social isolation versus males that were housed in groups with other males.

### Social isolation stress exacerbates the spread of TDP-43 protein pathology from surface glial cells to adjacent neurons

Because early life social isolation exacerbates the effects on lifespan of TDP-43 pathology induced only in SPG, we next tested the effects on the intercellular propagation of TDP-43 pathology to neurons. We previously have demonstrated that such focally induced TDP-43 protein pathology also results in non-cell-autonomous effects that include accumulation of TDP-43 pathology in nearby neurons that express physiological levels of human TDP-43 [34]. This propagation of toxicity between cell types is thought to play a major role in the progression of disease [14, 78–83]. In order to detect and quantify this intercellular effect, we used a humanized fly strain in which the endogenous TDP-43 homolog (TBPH) has been replaced by a human TDP-43 cDNA with a 3xFlag tag fused at the N-terminal (hTDP-43^WT-KI^). In this strain, the human TDP-43 functionally replaces the fly gene and is expressed under control of the fly ortholog’s promoter at physiological levels of hTDP-43 [34, 84]. On this background, we induced over-expression of hTDP-43 in the SPG. This system allows us to use phospho-specific antibodies to detect hyperphosphorylated, dysfunctional human TDP-43 within the SPG, where aggregation is induced by over-expression. But using this approach, we also detect the appearance of hyperphosphorylated, pathological TDP-43 protein in nearby neurons and glia that express physiological levels of the human TDP-43 protein [34]. This ‘spreading effect’ is progressive, with larger numbers of neurons exhibiting TDP-43 pathology and at increasing distance from the SPG over time.

We used the above system to test for impacts of social isolation stress on non-cell autonomous effects of glial hTDP-43 pathology on neurons. Because the impact on lifespan is relatively robust when male flies are isolated for 7 days (**Fig 1F**), we used that length of social isolation for the remainder of the experiments described. We quantified both the percentage of neurons that exhibit pathological human pTDP-43 and the distance of those neurons from the SPG. The neuronal nuclei were marked using an antibody against the pan-neuronal Elav marker. We visualized pTDP-43 using an established antibody that detects hyperphosphorylated TDP-43 protein [34] (**Fig 2**). In control animals that contain the hTDP-43^WT-KI^ but do not express the UAS-TDP-43 transgene in SPG, we observe no detectable pTDP43 signal at either 7 or 14 days after the temperature shift, irrespective of whether they had been previously group housed or isolated (**Fig 2 B,C, F, G**). By contrast we found, as previously reported, that induction of ectopic hTDP-43 in SPG was sufficient to cause accumulation of pTDP-43 in both the nucleus and cytoplasm of the SPG themselves, as well as in neurons. This effect of TDP-43 pathology in SPG on neuronal TDP-43 pathology was seen at both 7 and 14 days after induction of TDP-43 in the SPG (**Fig 2**). For instance, at just 7 days after inducing TDP-43 in SPG, we observe a small but significant number of neurons labelled with pTDP-43 (**Fig 2 D,J**), with increasing numbers of neurons labelled at 14 days post TDP-43 induction (**Fig 2 H,L**). The distance from the source of pathology in SPG also increased between 7 and 14 days after induction (**Fig 2 K,M**) as previously reported [34].

**Fig 2:**
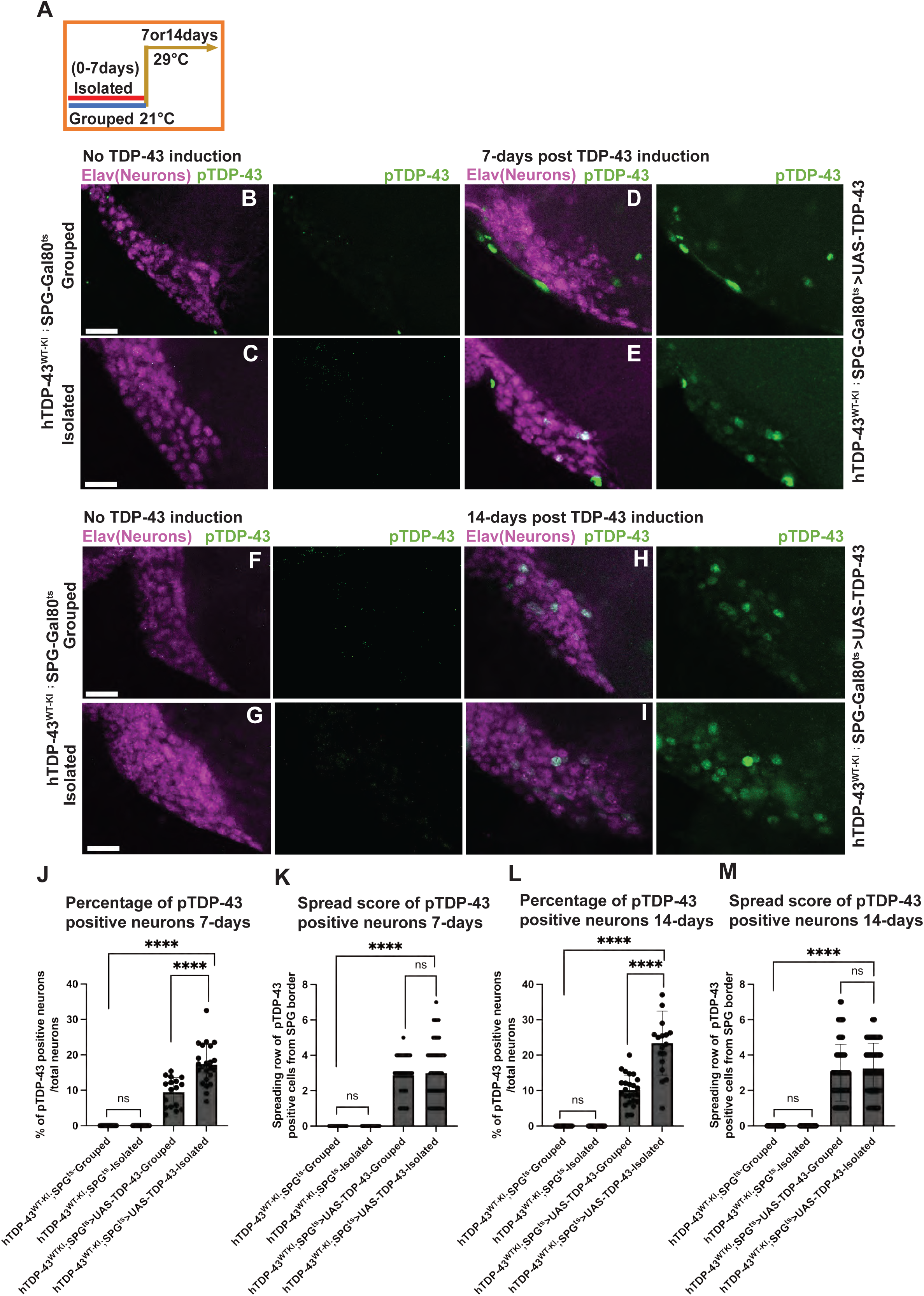
Social isolation prior to TDP-43 pathology induction causes a more rapid propagation of TDP-43 protein pathology from surface glial cells to adjacent neurons. For these experiments, a humanized fly model was used in which the endogenous fly ortholog (TBPH) is replaced with a functional copy of a sequence encoding human TDP-43 protein. The induced over-expression of human TDP-43 is restricted to SPG, but pathological effects in SPG trigger accumulation of pathologically hyper-phosphorylated TDP-43 in adjacent neurons. **A)** Schematic depicting 7-day early life 7-days social isolation or group housing of male flies at 21°C prior to TDP-43 induction in SPG followed by 7 or 14-days post TDP-43 induction in SPG post temperature shift induction at 29°C (B-M). (**B, C, J, K)** Immunofluorescence labeling indicates that without TDP-43 induction, there is no detectable pTDP-43 accumulation irrespective of whether the animals were grouped or isolated during the first 7 days of adult life. Scale bars= 50µm. hTDP-43^WT-KI^;SPG^ts^-Grouped(n=18),hTDP-43^WT-KI^;SPG^ts^-Isolated(n=18),hTDP-43^WT-KI^;SPG^ts^-Grouped (n=58), hTDP-43^WT-KI^;SPG^ts^-Isolated(n=58). Data shown are mean ± SEM and one-way ANOVA with Tukey’s multiple comparison test was performed ns:not significant. (**D)** By contrast, 7-days after TDP-43 over-expression in SPG (in the humanized hTDP-43 ^WT-KI^ background), there is already some pTDP-43 accumulation (as detected by pTDP-43 antibody; Green) in nearby neurons in grouped flies. (**E)** This effect is exacerbated by prior social isolation, leading to a significant increase in the percentage (**J**) of pTDP-43 labeled neurons (as detected by pTDP-43 antibody; Green). hTDP-43 ^WT-KI^;SPG^ts^>UAS-TDP-43-Grouped (n=17),hTDP-43^WT-KI^;SPG^ts^>UAS-TDP-43-Isolated (n=23). Data shown are mean ± SEM and one-way ANOVA with Tukey’s multiple comparison test was performed ****p<0.0001; but without changing the distance (**K**) of labeled neurons from the SPG source of TDP-43 pathology as seen in the quantification plots of spread score of pTDP-43 positive cells-7days. Neuronal nuclei are marked with the pan-neuronal Elav antibody (purple). Scale bars= 50µm. hTDP-43 ^WT-KI^;SPG^ts^>UAS-TDP-43-Grouped(n=75),hTDP-43^WT-KI^;SPG^ts^>UAS-TDP-43-Isolated (n=123). ns:not significant (**F and G)** In the absence of TDP-43 induction, pTDP-43 also is not detected 14 days after the temperature shift to 29°C, irrespective of whether the flies had previously been group housed or isolated during the first 7 days of adult life. Scale bar= 50µm. hTDP-43 ^WT-KI^;SPG^ts^-Grouped (n=15), hTDP-43 ^WT-KI^;SPG^ts^-Isolated (n=15), hTDP-43^WT-KI^;SPG^ts^-Grouped (n=29), hTDP-43^WT-KI^;SPG^ts^-Isolated(n=29), Data shown are mean ± SEM and one-way ANOVA with Tukey’s multiple comparison tests performed, ns:not significant. (**H,I)** By contrast, after 14-days of TDP-43 induction, we see a significant increase in the numbers of pTDP-43 positive neurons (as detected via pTDP-43 antibody; Green).But the numbers of neurons that exhibit such TDP-43 pathology are even higher following social isolation (**I,L**) vs group housing (**H, L**). Scale bars= 50µm. hTDP-43 ^WT-KI^;SPG^ts^>UAS-TDP-43-Grouped(n=24), hTDP-43 ^WT-KI^;SPG^ts^>UAS-TDP-43-Isolated (n=18). Data shown are mean ± SEM and one-way ANOVA with Tukey’s multiple comparison tests performed. ****p<0.0001. Here too, the average distance of labeled neurons from the SPG source of pTDP-43 is unchanged (**M**) as indicated in the spread score of positive pTDP-43 cells-14days. hTDP-43 ^WT-KI^;SPG^ts^>UAS-TDP-43-Grouped (n=74), hTDP-43^WT-KI^;SPG^ts^>UAS-TDP-43-Isolated (n=90). Data shown are mean ± SEM and one-way ANOVA with Tukey’s multiple comparison tests performed. ns:not significant.

We next examined the effects on neuronal TDP-43 pathology from social isolation stress. We found that isolating hTDP-43 ^WT-KI^;SPG-Gal80^ts^/UAS-TDP-43 male flies for 7 days prior to induction of TDP-43 results in a significantly higher percentage of neurons being labelled with pTDP-43 compared with animals of the same genotype that were group housed **(Fig 2)**. This increase in neuronal pathology from prior social isolation was evident at 4 (**S2 E Fig**), 7 and 14 days after TDP-43 induction in the SPG (**Fig 2 E,I**). But in contrast to the effects of social isolation on the number of neurons labelled (**Fig 2 J,L**) with pathological pTDP-43, the average distance of such labelled neurons from the SPG glia was not different between the grouped and isolated animals (**Fig 2 K,M**). These results demonstrate that prior social isolation stress exacerbates the inter-cellular effects of TDP-43 protein pathology in SPG that lead to pathology in nearby neurons. But the effect is to increase the total percentage of impacted neurons, without impacting the average distance of the impacted neurons from the source of toxicity in the SPG. The effect of social isolation thus appears to mainly impact the susceptibility of neurons to exhibit pathological changes in TDP-43 rather than the mechanism of intercellular effects of glia to neurons. These effects of social isolation on the number of neurons that exhibit TDP-43 protein pathology parallels the effects on lifespan in response to TDP-43 induction in SPG in SPG-Gal80^ts^/UAS-TDP-43 animals (**Fig 1**) as well as in the hTDP-43^WT-KI^;SPG-Gal80^ts^/UAS-TDP-43 animals that contain the human TDP-43 knocked in to replace the fly ortholog **(S2G Fig**).

### Social Isolation increases *Drosophila* mdg4-ERV activation in response to TDP-43 pathology

Work in a variety of animal models, cell culture and postmortem tissue has established that dysfunctional expression of endogenous retroviruses (ERVs) and retrotransposons (RTEs) is a hallmark of TDP-43 protein pathology [27–40]. We have previously demonstrated that the induction of pathological expression of TDP-43 in *Drosophila* glia, including SPG, is sufficient to cause expression and even replication of the mdg4-ERV [32–34]. The expression of this ERV also plays a key role in mediating both the intracellular and intercellular toxicity of glial TDP-43. We therefore examined the impact of social isolation on activation of mdg4-ERV.

We used an established mdg4-ERV reporter system, cellular labeling of endogenous retroviruses replication (CLEVR), that expresses nuclear mCherry upon mdg4 transposition [85]. We first quantified the effects of social isolation on the number of SPG that were labelled with the mdg4-CLEVR reporter in SPG-Gal80^ts^ flies that do not contain the TDP-43 transgene. As expected, there are very few SPG that exhibit mdg4-CLEVR reporter expression at either 7, 10 or 12 days after a temperature shift (**Fig 3 B, C, F, G and S3 B and C Fig**). By contrast, we see a significant increase in the number of mCherry labelled SPG nuclei after TDP-43 induction in SPG-Gal80^ts^/UAS-TDP-43 animals at each timepoint (**Fig 3 D, H and S3 D**), consistent with our previous report of mdg4 replication in response to TDP-43 protein pathology[33]. But we also see a dramatic increase in the number of mdg4-CLEVR positive nuclei in SPG-Gal80^ts^/UAS-TDP-43 animals that were previously stressed by social isolation for 7 days compared to those that were group housed prior to the TDP-43 induction. This effect was significant at both 10 and 12 days post TDP-43 induction (**Fig 3 E, I, J, and K**). At an earlier timepoint (7-days after TDP-43 induction) there are fewer SPG labeled with the mdg4-CLEVR reporter and no significant difference between the grouped vs isolated animals (**S3 Fig**). Together, these results indicate that the social isolation stress primes animals so that they exhibit higher levels of mdg4-ERV replication in response to subsequent induction of pathological TDP-43 levels in SPG.

**Fig 3:**
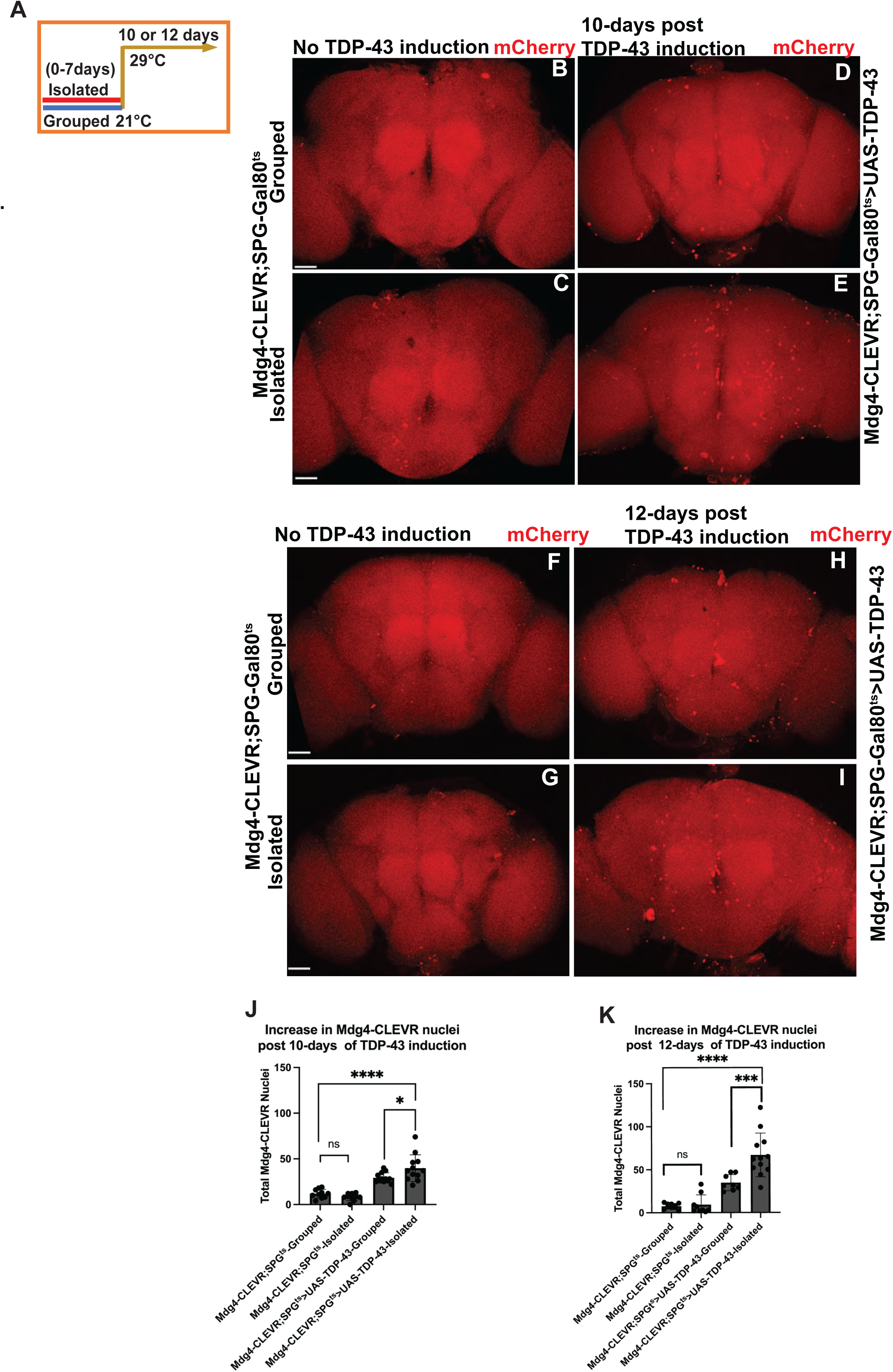
Early life social isolation causes more rapid mdg4-ERV replication in flies in response to TDP-43 induction in SPG. (**A)** Schematic illustrating isolated or grouped housing conditions for 7-days at 21°C prior to TDP-43 protein pathology induction and 10 or 12 days of TDP-43 induction after shifting to 29°C. (**B, C, J)** Quantification of mdg4-ERV reporter replication. We used cellular labeling of endogenous retroviruses replication (CLEVR), that expresses nuclear mCherry after mdg4 retrotransposition, to label glia in which the ERV replicates. Immunofluorescence images show that there is minimal mdg4-ERV replication in both grouped and isolated control flies without TDP-43 induction. Scale bars= 80µm. Mdg4-CLEVR; SPG^ts^-Grouped (n=10), Mdg4-CLEVR;SPG^ts^-Isolated (n=10); Data shown are mean ± SEM and one-way ANOVA with Tukey’s multiple comparison tests performed ns:not significant. (**D,E)** After 10 days of TDP-43 induction in SPG, we see SPG glial cells that exhibit reporter expression, indicating mdg4-ERV replication (as detected by mCherry, red). But there is a significant increase in the number of mdg4-CLEVR labelled SPG following isolation (**E,J**) compared to grouped housing (**D,J**) as indicated in the quantification plots of total number of mdg4-CLEVR nuclei post 10 days of TDP-43 induction. Scale bars= 80µm. Mdg4-CLEVR;SPG^ts^>UAS-TDP-43-Grouped (n=12), Mdg4-CLEVR; SPG^ts^>UAS-TDP-43-Isolated (n=12). Data shown are mean ± SEM and one-way ANOVA with Tukey’s multiple comparison tests performed. *p<0.05. (**F, G, K)** In the absence of TDP-43 expression, there is minimal mdg4-CLEVR reporter expression detected in both grouped and isolated flies at 12 days post temperature shift. Scale bars= 80µm. Mdg4-CLEVR; SPG^ts^-Grouped (n=10), Mdg4-CLEVR; SPG^ts^-Isolated (n=9); Data shown are mean ± SEM and one-way ANOVA with Tukey’s multiple comparison tests performed. ns: not significant. (**H, I)** By contrast, 12 days after TDP-43 induction we see many SPG labeled with the mdg4-CLEVR reporter (Cherry, red). But there is a significantly higher number of mdg4-CLEVR reporter labeled SPG after isolation **(I, K)** compared with grouped housing **(H, K)**. Scale bars= 80µm. Mdg4-CLEVR;SPG^ts^>UAS-TDP-43-Grouped (n=8), Mdg4-CLEVR; SPG^ts^>UAS-TDP-43-Isolated (n=12). Data shown are mean ± SEM and one-way ANOVA with Tukey’s multiple comparison tests performed. ***p<0.001.

### Flies need a holistic, enriched environment to provide protective effects of social interaction

Flies are known to socially interact during foraging, feeding, aggression, and courtship behaviors, and rely on group effects to protect from predators [63, 64, 74–77, 86–89].Social sensing is perceived via multiple sensory modalities that include visual, olfactory, and tactile perception [89–94]. To investigate the sensory requirements for the protective effects of group housing, we utilized a social divider tool with a mesh window that provided an intermediary between social isolation and social enrichment [95]. This transparent divider acted as a social barrier by dividing a food vial with an isolated fly on one side of the divider and 19 grouped flies on the other side (**Fig 4A**). The mesh window on the divider consists of holes with a pore size of 800 µm, sufficient to allow animals to extend legs to contact conspecifics on the other side of the barrier (As documented by video analyses,[95]). These dividers also should allow visual and olfactory contact between the isolated fly and the group on the other side.

**Fig 4:**
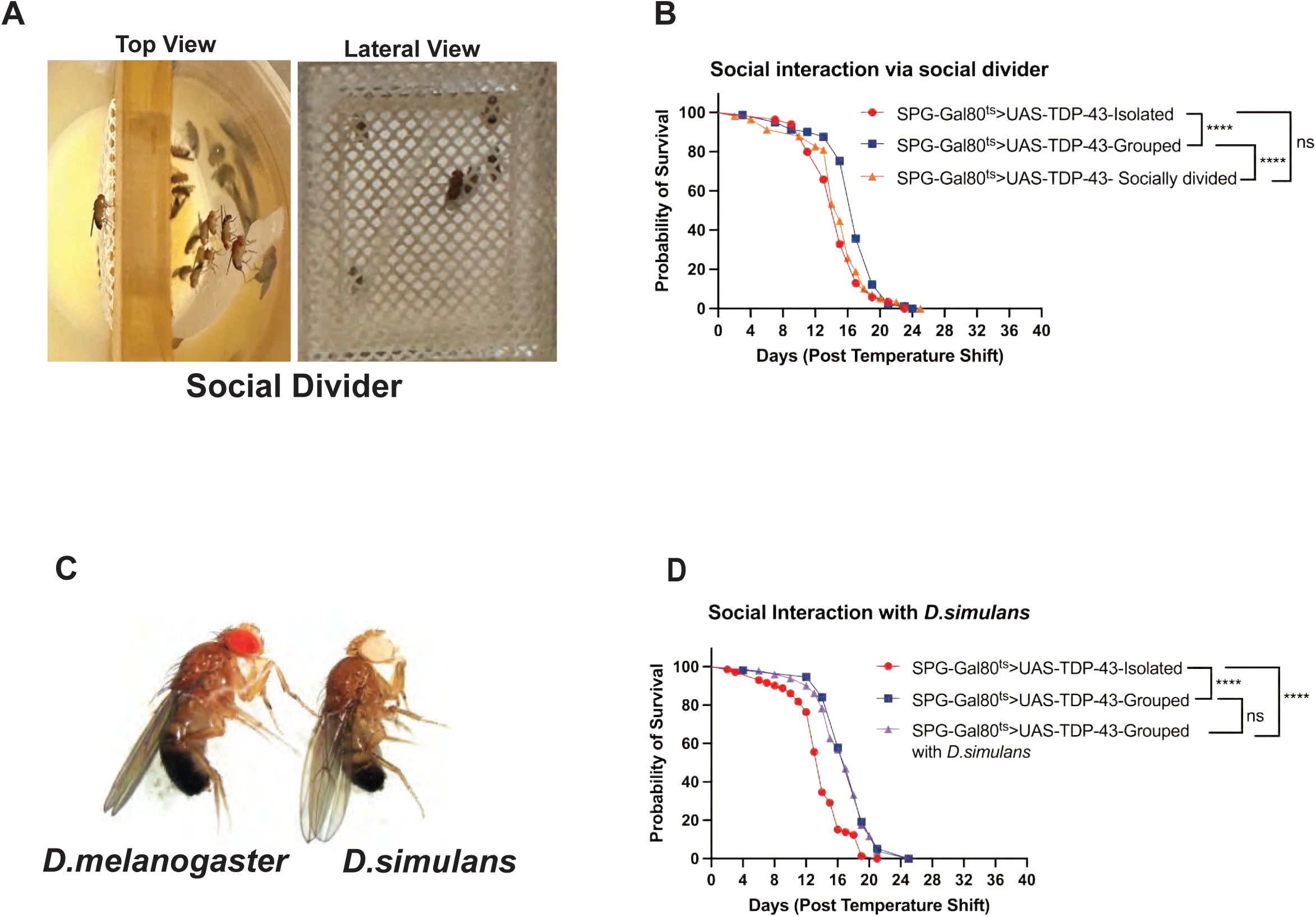
Flies require a holistically complete social interaction to protect from effects of social isolation. **A)** A social divider tool that allows visual, olfactory and tactile interaction, top and lateral views. Individual socially isolated flies were separated from a group of flies by placing a transparent divider with a mesh in the center of the divider with holes between them. **B)** Social interaction through the divider is not sufficient to provide the protective effects of group housing. SPG-Gal80^ts^>UAS-TDP-43 were either isolated, group housed, or allowed interaction only through the divider for their first 7 days of adult life. TDP-43 was then induced by shifting to 29°C and lifespans were measured in groups of 10. Median survival for the isolated flies was 15 (n=85), median survival of grouped flies was 17 (n=81), median life span of social divided flies was 15 (n=58). ****p< 0.0001 (Log-rank test). **C)** *D.melanogaster* vs *D.simulans* males are indistinguishable to the experimenter, except for the white eye color marker in the *D.simulans* stock. (**D)** Survival plots show that heterospecific social interaction of *D. melanogaster* males with *D.simulans* males is sufficient to rescue the effects of isolation on neurodegeneration. Survival curves are shown for SPG-Gal80^ts^>UAS-TDP-43 that were isolated, grouped with other melanogaster males or grouped with simulans males for the first 7 days of adult life. All groups were shifted to 29°C for TDP-43 induction after this 7 day period. Median survival for the isolated flies was 14 (n=72), median survival for males grouped with other melanogaster males was 19 (n=57), median survival for male *D.melanogaster* that had social interaction with *D.simulans* males was 17 (n=51). ****p< 0.0001 (Log-rank test) and ns:not significant. Kaplan-Meier – Simple survival analyses were performed for all survival plots.

We tested whether such partial social interaction through the social divider was sufficient to rescue the detrimental effects of isolation stress on subsequently induced TDP-43 pathology. We found that such partial interaction was not sufficient to rescue the social isolation effect, as the divided flies still experienced a significant decrease in survival rate when compared to the control grouped flies (**Fig 4B**). In fact, upon TDP-43 induction, the survival rate of divided flies does not differ from that of the fully isolated animals. This finding suggests that complete, more holistic social engagement with other individuals is needed to provide resilience to neurodegeneration later in life. Because these experiments involve only male flies, the beneficial impact of social interaction does not involve the rewards of mating. We also wondered whether the effects required interaction with a conspecific, or whether heterospecific social engagement, with males from a related species would suffice. To test this, we compared the effects of individual *Drosophila melanogaster* males housed alone, grouped with *D. simulans* males, or grouped with other *D. melanogaster* males. Because *D. simulans* and *D. melanogaster* have such similar size and physical appearance, we used white eye mutant *D. simulans* to aid in our own identification of the *D. melanogaster* males (**Fig 4C**). As previously, these grouped or isolated conditions were maintained for 7 days, followed by inducing SPG-specific expression of TDP-43 in the SPG-Gal80^ts^/UAS-TDP-43 target *D. melanogaster* males. Lifespans following induction were again measured in groups of 10. We found that social enrichment by housing *D. melanogaster* with either *D. melanogaster* or *D. simulans* to be equally protective against the effects of subsequent neurodegeneration. Indeed, the median survival in response to SPG-driven TDP-43 was equivalent among *D. melanogaster* that had previously been grouped with *D. simulans* or with D. melanogaster, while the isolated groups exhibited significantly reduced lifespans (**Fig 4D**).

Previous work has established that chronic social isolation stress leads to sleep loss in flies [9]. But reduced sleep is unlikely to be a primary driver of these effects for several reasons. First, although we do observe that social isolation reduces total and daytime sleep **(S4 A,B Fig)** as previously reported [9], we also found that treatment with gaboxadol, a GABA_A_ receptor agonist that is an established sleep aid in this model system [96] was not sufficient to ameliorate the effects of social isolation. Gaboxadol treatment did significantly increase both daytime and total sleep amount in both isolated and group housed animals **(S4 A, B Fig)**. Importantly, the levels of daytime and overall sleep levels in isolated flies treated with gaboxadol was higher than levels of daytime or overall sleep-in vehicle treated group housed flies. Despite this, such gaboxadol treatment was not insufficient to rescue the effects of social isolation on neurodegeneration (although curiously, gaboxadol treatment of group housed animals slightly reduced survival compared with vehicle treated group housed animals) **(S4 C Fig)**. We also tested the effects of severe sleep loss in *inc^1^*mutants, which sleep ∼60% less time than wild type control animals [97, 98]. on the rate of neurodegeneration. We found that the presence of the *inc^1^* mutation in hemizygous males had no effect on the length of lifespan after SPG-TDP-43 induction in group housed flies (**S4 D Fig**), indicating that even severe reductions of sleep that are observed in these mutants does not suffice to recapitulate the effects of social isolation stress. Interestingly, socially isolated and group housed *inc^1^* mutants do not differ in their lifespan responses to TDP-43 induction, exhibiting lifespans that are indistinguishable from that of group housed *inc+* animals. Thus, a genetic manipulation to reduce sleep levels is insufficient to sensitize group housed animals to neurodegeneration and in fact makes isolated animals resistant to those effects. Taken together, these findings indicate that reduced sleep is unlikely to be a major driver of the effects of social isolation on neurodegeneration.

## Discussion

Together, our findings indicate that early life social isolation stress accelerates the rate of progression of subsequently induced neurodegeneration in a relatively mild genetic model of TDP-43 dependent neurodegeneration in which the protein pathology is induced in just the SPG cell subtype. Flies are increasingly recognized as social animals, with multiple ethologically relevant benefits to group living that include access to mates, safety from predation [77, 88, 89, 93, 94, 99, 100], group foraging [101, 102] and the adaptive effects of laying eggs near the eggs of conspecifics [103–106]. In our assays, the impacts on sensitivity to the toxicity of TDP-43 pathology, loss of access to mating is not likely to be a stressor that contributes because group housing of males with other males is sufficient for the observed protective effects. In fact, even heterospecific interaction of *melanogaster* males with *simulans* males is protective. Because these closely related species share male pheromonal chemical cues [107–110], it is possible that chemical communication via species specific cuticular hydrocarbons plays a role in conveying awareness of the presence other individuals in our assay. But the protective effects of social interaction cannot be mediated by physical contact with other animals through a mesh divider. Rather, it requires that the animals occupy a shared physical space. This is consistent with the idea that a more multi-modal, holistic social interaction is required to protect animals from the stressful effects of isolation. These requirements for a wholistic social interaction to protect against subsequent neurodegeneration are similar to those that are required to protect from disruptive effects on sleep [95]. The findings reported here support the idea that chronic social isolation causes physiological stress that primes neurons and glial cells to be more susceptible to the toxicity of TDP-43 in a model of neurodegenerative disease.

Neurodegenerative diseases such as ALS, FTD and AD are thought to be triggered by a combination of genetic and environmental factors as well as the normal effects of aging. Epidemiological studies indicate that patients who suffer from neuropsychiatric disorders such as anxiety and depression or post-traumatic stress disorder, are at elevated risk of neurodegenerative diseases [1–3].The stress from social isolation also is a risk factor for these neurodegenerative disorders [4–8, 10–12]. Although the mechanisms remain largely unknown, studies from human and rodent models also indicate that a various types of early life adversity increase risk or severity of neurodegenerative diseases such as Alzheimer’s disease (AD) later in life [111]. But despite these strong correlative relationships, it is difficult to establish in human studies whether there are causative impacts on neurodegeneration from neuropsychiatric disorders, emotional stressors and other neurobehaviorally relevant aspects of life history. Animal models provide a means to investigate such causal relationships through controlled manipulations of life-history on the risk of onset, severity and rate of progression of neurodegeneration. Our identification of chronic social isolation stress in flies as an early life primer of TDP-43 toxicity establishes a platform to investigate the effects of life-history on risk of neurodegenerative disease.

## Supporting information

**S1 Fig:**
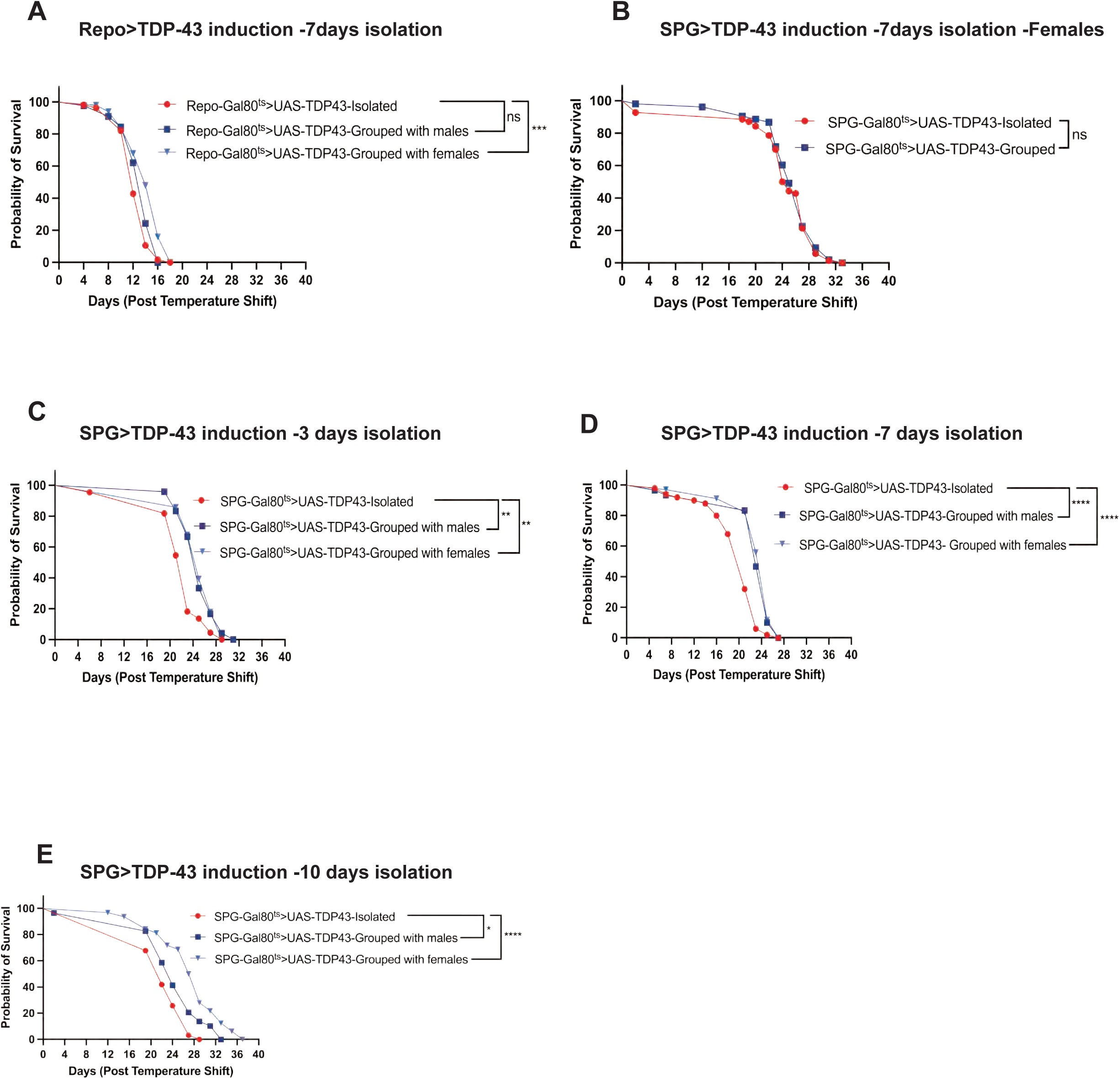
For male animals, social interaction with other males or with females are each protective against subsequent neurodegeneration. **A)** Survival curves are shown for male Repo-Gal80^ts^>UAS-TDP-43 flies after induction of TDP-43 in all glia. Such males were either socially isolated for the first 7 days of adult life (Isolated) or they were housed with 19 other males (Grouped with males) or they were housed with 19 females (Grouped with females). After this 7 day period at 21°C, the temperature was shifted to 29°C for induction of TDP-43 in all glia. The effects of social isolation vs group housing are less clear here compared with the context of more limited expression in just the SPG glia. This likely has to do with the severity of the effects of such pan-glial expression which results in an extremely shortened lifespan. In this context, social isolation has no effect compared to group housing with other males, and has a modest but significant effect compared to group housing with females. Median survival of male flies after social isolation was 12 (n=56), vs 14 (n=45) for males that were group housed with other males prior to induction. Median life span of males that were group housed with females prior to TDP-43 induction was 14 (n=50). ***p<0.001, ns:not significant (Log-rank test). **B)** 7-day social isolation has no effects on survival in females after subsequent TDP-43 induction in SPG. Survival curves are shown for SPG-Gal80^ts^>UAS-TDP-43. Median life span of isolated females was 24.5 (n=70), median life span of grouped females was 25 (n=53). ns:not significant (Log-rank test). (**C,D,E)** For male flies, effects on later neurodegeneration from early life social interaction with either other males or with other females are largely equivalent. Males were isolated, grouped with other males or group housed with females for the first 3 (**C**), 7 (**D**) or 10 (**E**) days of adult life. Median survival of males after TDP-43 induction following 3-days of social isolation was 23 (n=22), compared with 25 (n=24) after 3 days of group housing with other males and 25 (n=28) when grouped with females. **p<0.01 (Log-rank test). **D)** Median survival with TDP-43 induction after 7-days isolation was 21 (n=50) compared to 23 (n=30) after group housing with males and 25 n= (34) after group housing with females. ****p<0.0001 (Log-rank test). **E)** Median survival with TDP-43 induction after 10-days of isolation was 22 (n=31), compared to 24 (n=29) after group housing with males and 28 n= (32) after group housing with females. *p<0.05; ****p<0.0001 (Log-rank test).

**S2 Fig:**
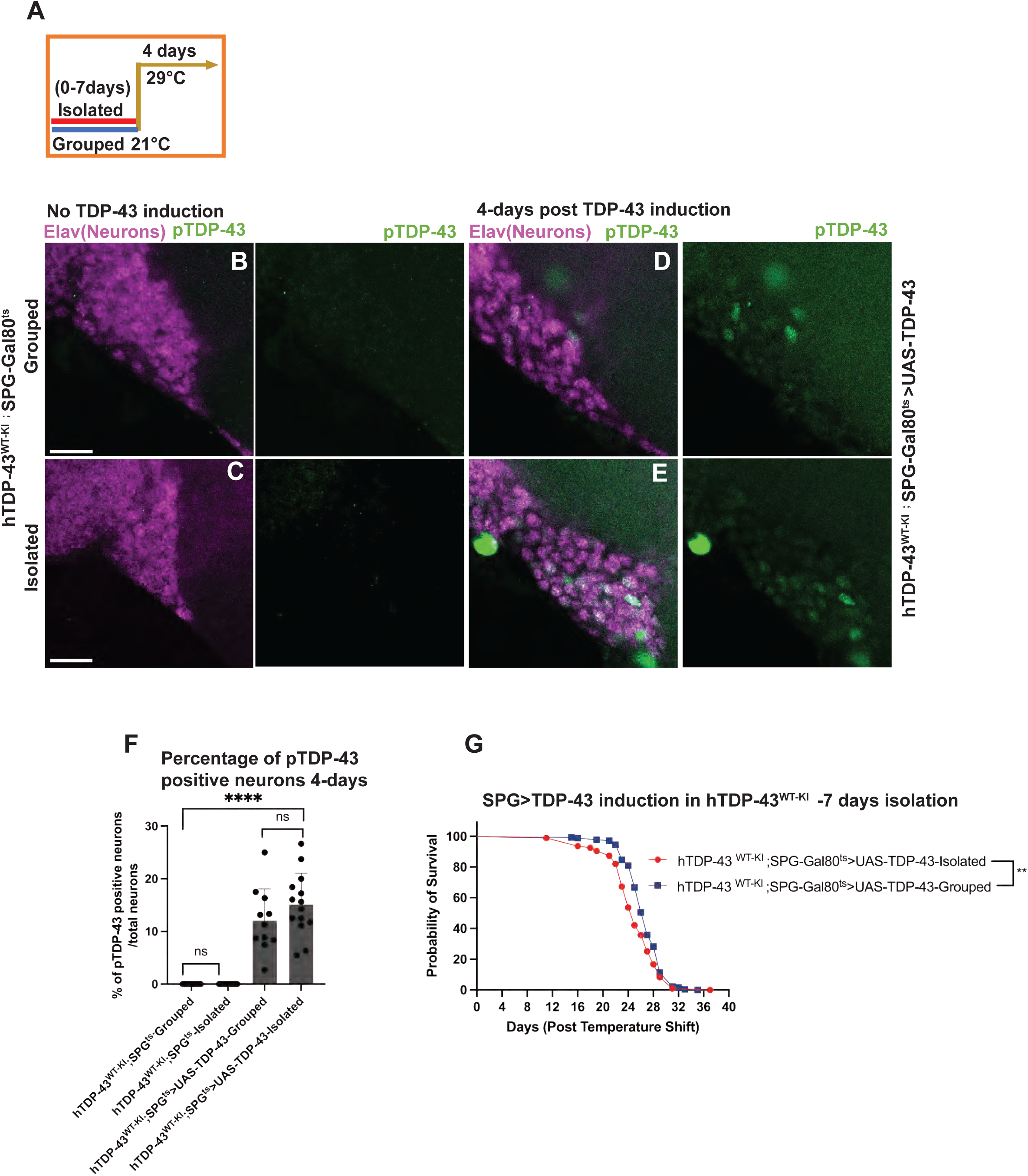
Social isolation stress prior to TDP-43 induction in SPG exacerbates the spread of TDP-43 pathology from surface glia to nearby neurons. **A)** Schematic represents 7-days social isolation vs group housing paradigm at 21°C prior to TDP-43 pathology induction for 4-days at 29°C. These experiments were done using hTDP-43 ^WT-KI^;SPG^ts^>UAS-TDP-43 animals that contain a humanized TDP-43 knocked in to replace the fly ortholog, and also contain an inducible human TDP-43 that is expressed in SPG after temperature shift. hTDP-43 ^WT-KI^;SPG^ts^ animals that contain the human knock in but do not over-express TDP-43 in SPG were used as a genetic control. In the genetic controls (**B, C, F)** observe no detectable accumulation of pTDP-43 after grouped or isolated housing. Scale bar= 50µm. hTDP-43 ^WT-KI^;SPG^ts^-Grouped (n=12,) hTDP-43 ^WT-KI^;SPG^ts^-Isolated (n=21). Data shown are mean ± SEM and one-way ANOVA with Tukey’s multiple comparison tests performed, ns:not significant. By contrast, after induction of TDP-43 in the SPG (**D, E, F)** we see significant accumulation of pTDP-43 in neurons nearby to the SPG. This is apparent both in the animals that were grouped or isolated prior to TDP-43 induction. At this timepoint (4 days after TDP-43 induction), there is no significant difference in the percentage of neurons that exhibit pTDP-43 accumulation (**F**). hTDP-43 ^WT-KI^;SPG^ts^>UAS-TDP-43-grouped (n=11), hTDP-43 ^WT-KI^;SPG^ts^>UAS-TDP-43-isolated (n=14). Data shown are mean ± SEM and one-way ANOVA with Tukey’s multiple comparison tests performed. ns:not significant. **G)** Significant effects of social isolation are seen on median survival in response to TDP-43 induction in hTDP-43^WT-KI^;SPG-Gal80^ts^/UAS-TDP-43 flies, as is seen with the same experiment done in animals that do not contain the human knock in (see main text). Survival curves are shown for hTDP-43^WT-KI^;SPG-Gal80^ts^>UAS-TDP-43. Median survival of hTDP-43^WT-KI^;SPG-Gal80^ts^>UAS-TDP-43 isolated flies was 25 (n=95), median survival of hTDP-43^WT-KI^;SPG-Gal80^ts^>UAS-TDP-43 grouped males was 27 (n=184).

**S3 Fig:**
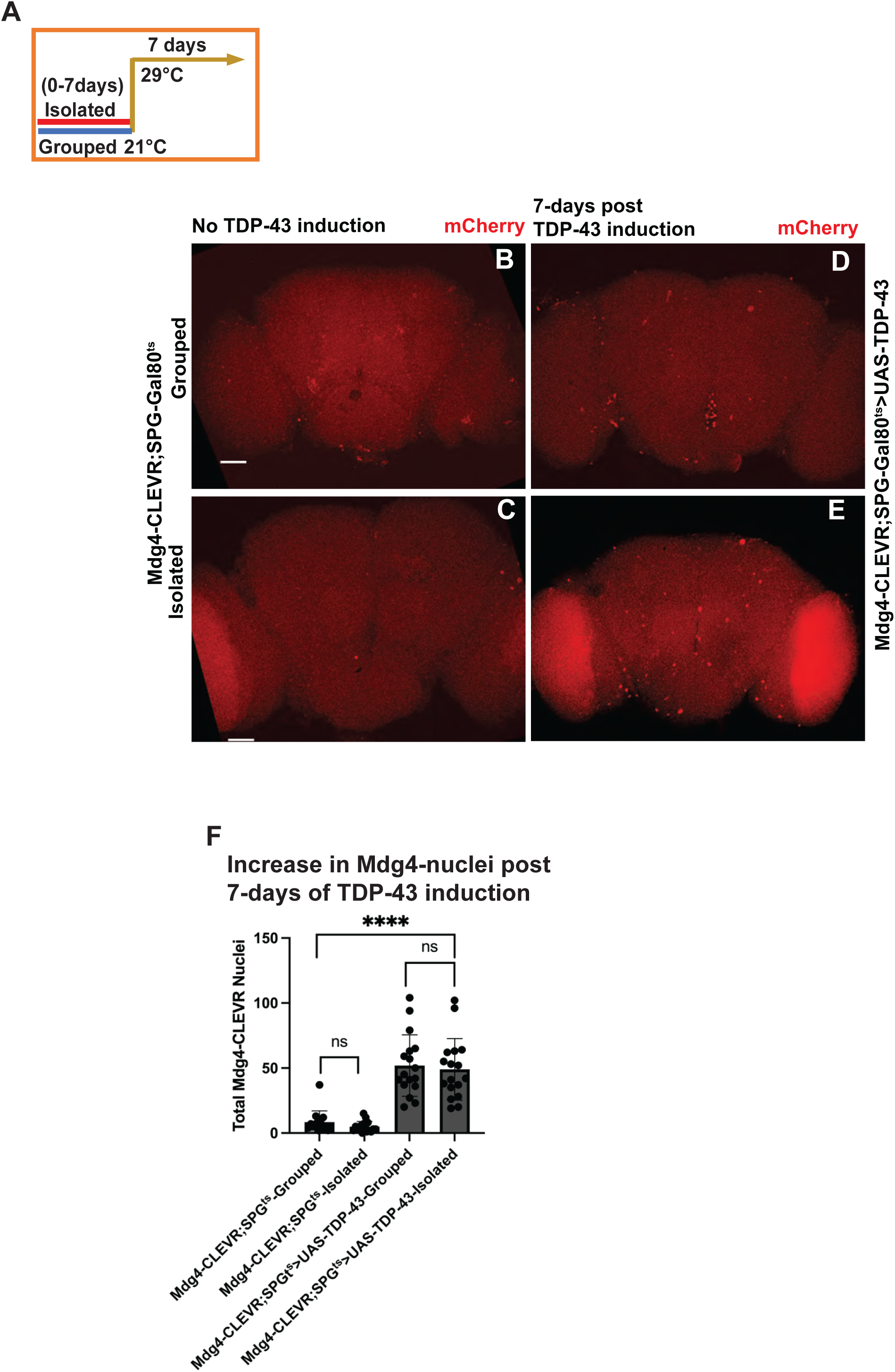
Early life social isolation stress causes rapid activation of Mdg4-ERV replication in flies in response to TDP-43 induction in SPGs (surface glia). **A)** Schematic indicates isolated or grouped housed conditions for 7-days at 21°C prior to TDP-43 induction and post 7-days of TDP-43 protein pathology induction after shifting to 29°C. (**B and C)** Few SPG are labeled with the mdg4-CLEVR reporter of ERV replication in grouped and isolated control flies without TDP-43 induction. Mdg4-CLEVR; SPG^ts^-Grouped (n=15), Mdg4-CLEVR;SPG^ts^-Isolated (n=17). Data shown are mean SEM and one-way ANOVA with Tukey’s multiple comparison tests performed. ns:not significant. (**D,E)** After 7 days induction of TDP-43 in SPG there are a significant number of mdg4-CLEVR labeled SPG both in animals that were grouped or isolated for the first 7-days of adult life, prior to TDP-43 induction in SPG. Scale bar= 80µm. But at this timepoint, 7 days after TDP-43 induction, there is no significant difference in the number of nuclei labelled with the mdg4-CLEVR mCherry reporter between the animals that were previously isolated or grouped (F). Mdg4-CLEVR;SPG^ts^>UAS-TDP-43-Grouped (n=17), Mdg4-CLEVR; SPG^ts^>UAS-TDP-43-Isolated (n=17). Data shown are mean SEM and one-way ANOVA with Tukey’s multiple comparison tests performed, ns:not significant.

**S4 Fig:**
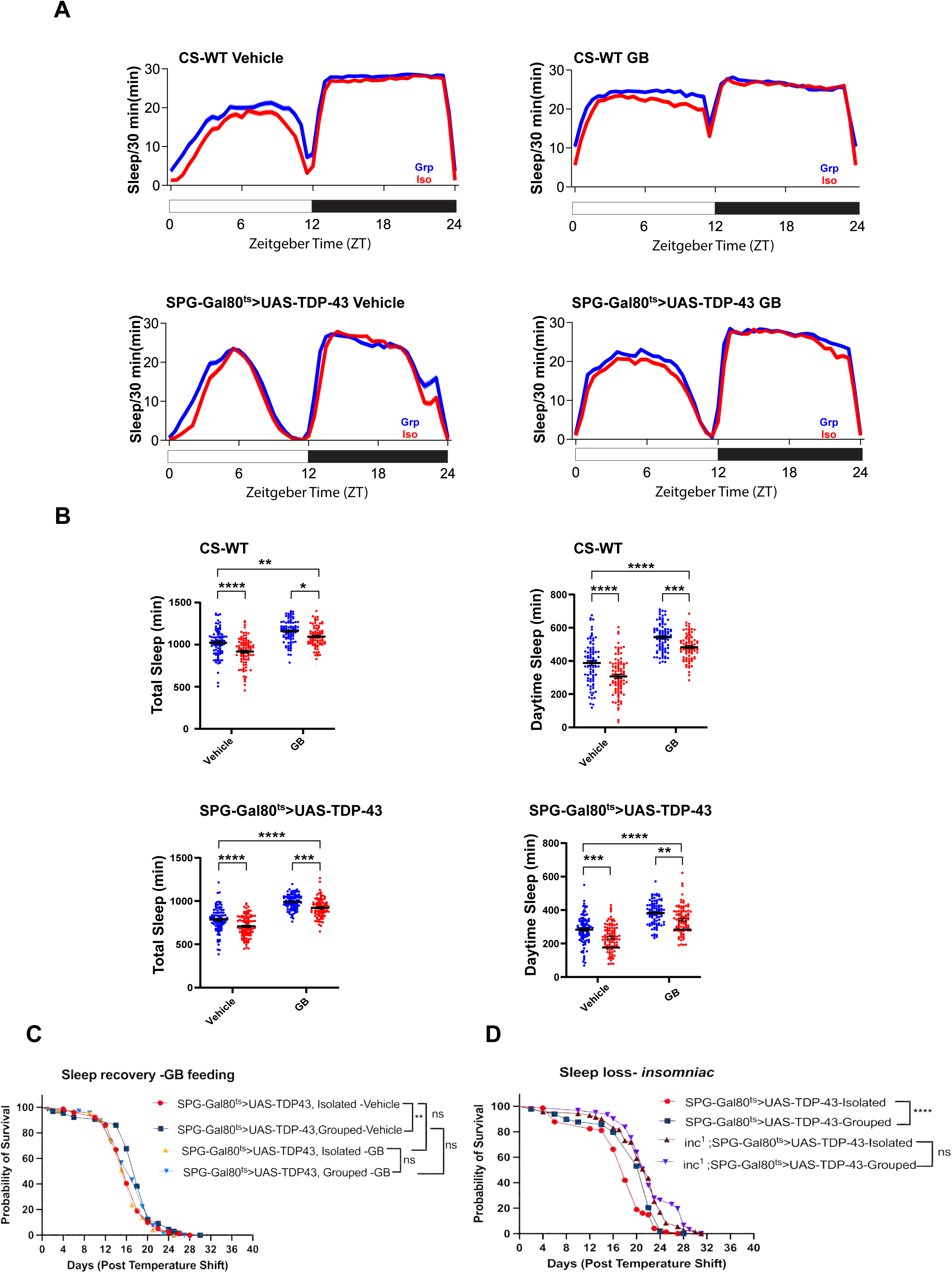
Sleep loss caused by social isolation is not the main driver of the effects on neurodegeneration from social isolation. (A) Social isolation of either CS-WT or SPG-Gal80ts>UAS-TDP-43 causes a reduction in both total and daytime sleep relative to that of group housed flies. Feeding either of these genotypes of flies with gaboxadol (GB), a GABA_A_ receptor agonist was sufficient to restore sleep in isolated flies to levels at or above that normally seen in group housed animals (A and B). Gaboxadol treated isolated flies exhibit levels of both daytime and total sleep that are as high or higher than that of grouped flies that are not treated with gaboxadol. (B) Quantification of sleep across each individual fly shown for the same genotypes/treatments and sample sizes as in (A).Sleep profiles (min/30 min) across a 24-hr light:dark cycle (ZT 0–24) for group-housed (Grp, blue) vs. isolated (Iso, red) flies CS-WT fed vehicle (Grp n=83, Iso n=92) or gaboxadol (GB) (Grp n=90, Iso n=84), SPG^ts^>UAS-TDP-43 fed vehicle (Grp n=95, Iso n=94) or GB (Grp n=95, Iso n=90). Two-way ANOVA (housing × treatment) with Šídák’s multiple comparisons test: Socal isolation significantly reduces sleep and treating the flies gaboxadol significantly increased sleep levels in all datasets (both factors p < 0.0001), with no significant interaction (p > 0.05), indicating additive, independent effects; *p < 0.05, **p < 0.01, ***p < 0.001, ****p < 0.0001; ns, not significant. Two-way ANOVA (housing× treatment) with Šídák’s multiple comparisons test: *p < 0.05, **p < 0.01, ***p < 0.001, ****p < 0.0001; ns, not significant **C)** Despite the fact that GB feeding restores sleep of isolated flies to levels at or above that of group housed flies not fed GB, this is not sufficient to ameliorate the effects of social isolation on neurodegeneration. Survival plots show that even with sleep levels restored during the 7-day isolation, the effects of isolation on lifespan after TDP-43 induction remains. Survival curves are shown for SPG-Gal80^ts^>UAS-TDP-43 genotypes. Median survival of SPG-Gal80^ts^>UAS-TDP-43, vehicle-Isolated was 16 (n=79), median survival of SPG-Gal80^ts^>UAS-TDP-43 vehicle-Grouped was 18 (n=65), median survival of gaboxadol treated SPG-Gal80^ts^>UAS-TDP-43-Isolated flies was 17 (n=71), median survival of gaboxadol treated SPG-Gal80^ts^>UAS-TDP-43-Grouped flies was 17 (n=67). **p< 0.01 (Log-rank test) and ns:not significant. **D)** Despite the known decrease in total sleep of insomniac *(inc1)* mutants, the survival of group housed inc1 mutants after TDP-43 induction in SPG was indistinguishable from that of group housed wild type flies, indicating that sleep loss per se is not sufficient to prime flies for faster neurodegenerative effects in response to SPG>TDP-43 induction. Median survival of SPG-Gal80^ts^>UAS-TDP-43 Isolated was 18 (n=74), median survival of SPG-Gal80^ts^>UAS-TDP-43 Grouped was 22 (n=49), median survival of inc^1^; SPG-Gal80^ts^>UAS-TDP-43-Isolated was 22 (n=71), inc^1^;SPG-Gal80^ts^>UAS-TDP-43-Grouped was 22 (n=61). ****p< 0.0001 (Log-rank test) and ns:not significant. Kaplan-Meier – Simple survival analyses were performed for all survival plots.

## Material and Methods

### Table for materials used in this study (Table1)

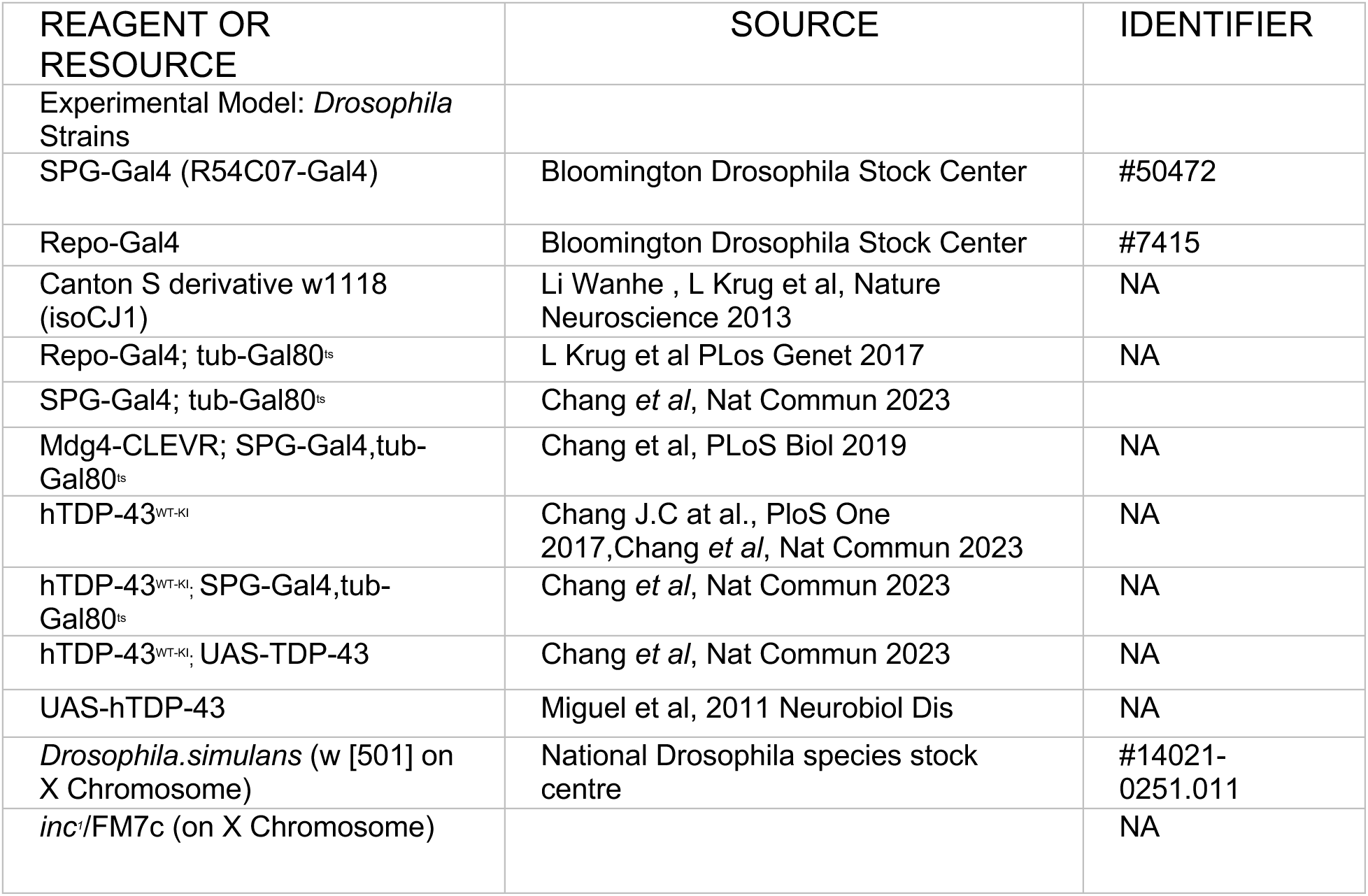

### Fly medium and stocks

Flies were raised on standard corn meal media at 25°C and 21°C incubators in a 12h light/dark (LD) cycle. A list of all the fly stocks used in this study are listed below in the resources table. All our stocks were back crossed with CantonS derivative *w^1118^* (isoCJ1) [34] our in-house wild type strain.

### Social isolation paradigm and survival assays

Flies were subjected to social isolation by placing a newly eclosed single male flies in a standard food vial, and for social enrichment, the flies were group-housed with other males or females as indicated in groups of 20 in a standard food vial. The flies were regrouped into cohorts of 10 to measure their lifespans. We used an inducible gene expression system-Gal80^ts^-for cell type specific overexpression with either pan glial driver line-Repo-Gal4, or subtype glial driver line-SPG-Gal4. The flies were grown at 21°C from embryonic stages for normal development. The 0-day old flies were isolated or grouped for a period of 1,3,5,7 or 10 days and there was no TDP-43 induction during isolation/grouped condition. These isolated and grouped flies were temperature shifted to 29°C for inducible TDP-43 expression via Gal80^ts^ and used for life span monitoring, and for brain dissections. The 7-day isolation paradigm was chosen for all the experiments including most of the life spans and brain dissections throughout the manuscript as the most robust effects were found from the 7-day isolation paradigm.

### Brain Dissections and Immunohistochemistry

Fly adult brains were dissected in ice-cold phosphate buffered solution (PBS). Dissected brains were quickly transferred into 4% paraformaldehyde-PFA (Electron Microscopy sciences #15713) for fixation diluted in PBS solution (1X PBS+4%PFA+0.2% Triton-X-100 (Sigma Aldrich)) and then incubated for a 30 mins interval under vacuum for 2 times. The fixated brains were washed four times at 15 min intervals with 1X PBST sample buffer (1X PBS+ 1%Triton-X-100+ 3%Nacl). Brain samples were then incubated in a blocking solution (1XPBST+10% normal horse serum (NHS) overnight at 4°C on a nutator. After blocking step, the brains were then incubated with primary antibodies diluted in 1X PBST at required dilutions overnight at 4°C. Samples were then washed with sample buffer solution (1X PBS+ 1%Triton-X-100+ 3%Nacl) for four times at 15 mins interval at room temperature on a nutator. Primary incubated brains were then transferred to secondary antibodies diluted in 1X PBST + 10% NHS and incubated overnight at 4°C.Primary and secondary antibodies used are listed below in the table. The brain samples were washed with a sample buffer solution again four times at 15 mins interval at room temperature on a nutator. The washed brain samples were then mounted on clean slides using focus clear mounting media (CelExplorer, FC-101). Mounted brain samples were then imaged using a Zeiss LSM 800 microscope, and images were processed using image J.

### Social divider assay

The social divider tool consisted of transparent divider that was placed in a food vial that created physical barrier between the isolated male fly and 19 grouped flies on the other side of the divider mesh. The divider included a mesh window in the center of the divider with a hole’s size of 800 µm that allows social interaction between the isolated and grouped flies by allowing the flies to smell, see and touch by placing their legs through the holes. This condition was sustained for 7 days at 21°C. After 7 days, the singular divided flies were regrouped into groups of ten, and their temperature was shifted to 29°C for TDP-43 induction and lifespan measurements.

### Social interaction assay with *D.simulans*

Social interaction between the *D. melanogaster* and its distant relative *D. simulans* was measured by placing isolated *D. melanogaster* with 19 grouped *D. simulans* in a food vial. The key distinguishing factor to differentiate between the two fly species was the eye color. The *D. melanogaster* flies had red eyes and *D. simulans* had white eyes.

### Social isolation paradigm for sleep assays

Newly eclosed flies were collected and kept in standard fly food bottles (∼200 flies per bottle) for 3–5 days for social enrichment and mating. Male flies were then distributed as single fly in a food vial for achieving social isolation (Iso), and 25 flies in a food vial for social enrichment (Grp).

### Sleep Assays

Sleep profiles of flies were monitored using the DAM system (TriKinetics, Waltham, MA). Flies were maintained in isolated or grouped conditions for 7 days, followed by a 3-day monitoring period of sleep. Sleep was measured as previously described[9].Flies were treated with gaboxadol drug both during isolation/grouped and sleep monitoring period. After treatment with gaboxadol, the total sleep time, daytime sleep and nighttime were measured.

### Gaboxadol Treatment

Gaboxadol hydrochloride (Sigma Aldrich-T101; CAS number: 85118-33-8) [96]was dissolved in water to make stock solution at a higher concentration of (1mg/ml). A final concentration of 0.01mg/ml of gaboxadol drug was dissolved in cooled and melted fly food for treatment of gaboxadol flies. Vehicles contained similar amounts of water dissolved in fly food. Both grouped and isolated flies were treated with gaboxadol for a duration of 7 days.

### Statistics

Statistical analyses for individual experiments were performed using Graphpad Prism 10. Specific statistical analyses conducted for showing significance of experimental groups and specific significance tests are mentioned in each of the individual figure legends. Survival assays were analyzed using the Kaplan-Meier method from Graphpad Prism. The significant levels were indicated as star numbers in the following format: *p<0.05, **p<0.01, ***p < 0.001 and ****p < 0.0001.

## Resource Avaliability

### Lead contact and material availability

Requests for resources and reagents should be directed to and will be fulfilled by the lead contact, Josh Dubnau. All Drosophila strains will be made available upon request to the lead contact.

### Data and code availability

This study did not generate or analyzed any datasets/code.

## Acknowledgements

We thank Yangyuan Li for help with making dividers. We thank members of Josh Dubnau lab, Roger Sher lab and Johnathan Nelson lab for comments and suggestions on the experiments.

## Author Roles

Swetha MurthyGowda : Conceptualization, Data curation, Formal analysis, Investigation, Writing -reviewing & editing

Kyle Huyghue: Data curation, Formal analysis, Investigation, Writing -reviewing & editing

Brissa Castillo: Data curation-performed sleep assays

Fiona Gugala: Data curation-performed sleep assays

Joshua Dubnau: Conceptualization, Funding acquisition, Investigation, Writing - reviewing & editing

